# Pupillary Dynamics of Mice Performing a Pavlovian Delay Conditioning Task Reflect Reward-Predictive Signals

**DOI:** 10.1101/2022.09.15.508189

**Authors:** Kota Yamada, Koji Toda

## Abstract

Pupils can signify various internal processes and states, such as attention, arousal, and working memory. Changes in pupil size have been associated with learning speed, prediction of future events, and deviations from the prediction in human studies. However, the detailed relationships between pupil size changes and prediction are unclear. We explored pupil size dynamics in mice performing a Pavlovian delay conditioning task. A head-fixed experimental setup combined with deep-learning-based image analysis enabled us to reduce spontaneous locomotor activity and to track the precise dynamics of pupil size of behaving mice. By setting up two experimental groups, one for which mice were able to predict reward in the Pavlovian delay conditioning task and the other for which mice were not, we demonstrated that the pupil size of mice is modulated by reward prediction and consumption, as well as body movements, but not by unpredicted reward delivery. Furthermore, we clarified that pupil size is still modulated by reward prediction even after the disruption of body movements by intraperitoneal injection of haloperidol, a dopamine D2 receptor antagonist. These results suggest that changes in pupil size reflect reward prediction signals. Thus, we provide important evidence to reconsider the neuronal circuit involved in computing reward prediction error. This integrative approach of behavioral analysis, image analysis, pupillometry, and pharmacological manipulation will pave the way for understanding the psychological and neurobiological mechanisms of reward prediction and the prediction errors essential to learning and behavior.

**Manuscript contributions to the field:** Predicting upcoming events is essential for the survival of many animals, including humans. Accumulating evidence suggests that pupillary responses reflect autonomic activity and are modulated by noradrenergic, cholinergic, and serotonergic neurotransmission. However, the relationships between pupillary responses, reward prediction, and reward prediction errors remain unclear. This study examined changes in pupil size while water-deprived mice performed a Pavlovian delay conditioning task using a head-fixed setup. The head-fixed experimental setup, combined with deep-learning-based image analysis, enabled us to reduce spontaneous locomotor activity and to track the precise dynamics of the licking response and the pupil size of behaving mice. A well-controlled, rigid behavioral experimental design allowed us to investigate the modulation of behavioral states induced by reward prediction. While pharmacological manipulation might affect pupil size, the combined approach of pupillometry and pharmacological manipulation allowed us to differentiate reward prediction signals and signals modulated by body movements. We revealed that the changes in pupil size (1) reflect reward prediction signals and (2) do not reflect signals of reward prediction error. These results provide novel insights into the neuronal circuitry potentially involved in computing reward prediction errors. The integrative approach of behavioral analysis, image analysis, pupillometry, and pharmacological manipulation used in this study will pave the way for understanding the psychological and neurobiological mechanisms of prediction and the prediction errors essential in learning and behavior.

## Introduction

Predicting future events from current observations helps organisms to obtain rewards and avoid aversive events in a given environment. Pavlovian conditioning is a widely used experimental procedure for investigating the predictive abilities of animals. For example, water-restricted mice are exposed to an auditory stimulus, followed by the water reward. After several training sessions, mice develop anticipatory responses to the auditory stimulus. Pavlovian conditioning involves both behavioral and physiological responses. In appetitive conditioning, a conditioned approach response to a stimulus that signals food (Hearst and Jenkins, 1974) or to the location where the food is presented (Boakes, 1977) is observed. In fear conditioning, freezing responses (Estes and Skinner, 1941) are induced by a stimulus that signals aversive events. Physiological responses, such as salivary response, changes in skin conductance, heart rate, pupil dilation, body temperature, and respiration, are also acquired through Pavlovian conditioning (Esteves et al., 1994; Leuchs et al., 2017; Lonsdorf et al., 2017; Notterman et al., 1952; Öhman et al., 1976; Ojala and Bach, 2020; Pavlov, 1927; Pietrock et al., 2019; Wood and Obrist, 1964). Pavlovian conditioning includes several response types: preparatory, consummatory, and opponent responses to unconditioned responses (Konorski, 1967; Solomon and Corbit, 1974). Thus, accumulating evidence in the field of psychological and physiological studies of animal learning demonstrates that Pavlovian conditioning is a valuable technique for studying the function and underlying mechanism of prediction.

Although the use of pupillometry in Pavlovian conditioning dates back more than half a century, its reliability as an indicator of learning has recently been reevaluated (Finke et al., 2021). It has been reported that changes in pupil size occur as a reactive response to a conditioned stimulus in fear and appetitive conditioning in humans (Leuchs et al., 2017; Lonsdorf et al., 2017; Ojala and Bach, 2020; Pietrock et al., 2019). The relationship between pupil size and theories of learning, such as prediction errors in temporal difference learning (Sutton and Barto, 2018), the Mackintosh and Rescorla-Wagner models (Mackintosh, 1975; Rescorla and Wagner, 1972), as well as attention to the stimuli in the Pearce-Hall model (Pearce and Hall, 1980) have also been discussed (Koenig et al., 2018; Pietrock et al., 2019; Vincent et al., 2019). Changes in pupil size are associated with various internal states, including arousal level, attention, working memory, social vigilance, the value of alternatives in choice tasks, and uncertainty in diverse research fields (Ebitz et al., 2014; Ebitz and Platt, 2015; Finke et al., 2021; Joshi and Gold, 2020; Larsen and Waters, 2018; Van Slooten et al., 2018; Vincent et al., 2019; Zénon, 2019). These findings suggest that pupil size is a reactive response to a conditioned stimulus and an active modulator of sensorimotor processing that affects prediction (Ebitz and Moore, 2019).

Despite the potential usefulness of pupillometry in understanding the neurobiological mechanisms underlying behavior, there have been only a few attempts to record pupillary changes in rodent research (Cazettes et al., 2021; Lee and Margolis, 2016; Nelson and Mooney, 2016; Privitera et al., 2020; Reimer et al., 2014; Wang et al., 2022). This can be attributed to two technical issues. First, conventional behavioral tasks designed for rodents use experimental apparatuses in which animals move freely, making it impossible to precisely record pupil size. Second, body movements also modulate pupil size (Cazettes et al., 2021; Nelson and Mooney, 2016). This makes its interpretations more complex than human studies that allow participants to remain in the experimental setup. Recent experimental setups and advances in machine learning have enabled researchers to overcome these technical limitations. By combining a head-fixed setup with image analysis techniques such as DeepLabCut (Mathis et al., 2018; Nath et al., 2019), several studies have successfully quantified pupils and eyelid size of mice performing behavioral tasks (Kaneko et al., 2022; Privitera et al., 2020).

This study explored the dynamics of licking and pupillary responses of mice performing a Pavlovian delay conditioning task with a head-fixed experimental setup. Pupil size is known to be increased by the presentation of the cue in appetitive and aversive conditioning in human participants as well as rodent subjects, supporting the view that animals gain their arousal by the cue presentation (Finke et al., 2021; Pietrock et al., 2019). In Experiment 1, we trained head-fixed mice on the Pavlovian delayed conditioning task. An auditory stimulus was presented before a sucrose solution reward was delivered while recording their licking and pupil response. In this task, we designed contingent and non-contingent groups to manipulate the predictability of the delivery of the sucrose solution by the auditory stimulus. In the contingent group, the auditory stimulus signaled the arrival of the sucrose solution, such that the delivery of the sucrose solution immediately followed the auditory stimulus. In the non-contingent group, the auditory stimulus provided no predictive information about the arrival of the sucrose solution, as the presentation of the auditory stimulus and the delivery of the sucrose solution were randomized. We investigated the dynamics of licking and pupillary responses in predictable and unpredictable situations by measuring licking and pupillary responses while the mice performed the Pavlovian delay conditioning tasks. In addition, bout analysis of the licking responses allowed us to unveil the detailed relationship between the licking and pupillary responses. In Experiment 2, we examined the pupil dynamics by suppressing body movements with systemic administration of haloperidol, an antagonist of dopamine D2 receptors that have been reported to inhibit anticipatory and consummatory licking (Fowler and Mortell, 1992; Liao and Ko, 1995) and spontaneous movements in an open-field experiment (Arruda et al., 2008; Bernardi et al., 1981; Conceição and Frussa-Filho, 1996; Strömbom, 1977).

## Methods

### Subjects

Eight adult male C57BL/6J mice were used. All mice were purchased from Nippon Bio-Supp.Center and bred in the breeding room provided in the laboratory. All mice were naive and eight weeks old at the start of the experiment. The mice were maintained on a 12:12 h light cycle. All the experiments were conducted during the dark phase of the light cycle. The mice had no access to water in their home cage and were provided with water only during experimental sessions. The mice were allowed to consume sufficient sucrose solution during the experiment. The mice’s body weight was monitored daily (21.7 ± 2.1 g before the experiment). They were provided additional access to water at their home cage if their body weight fell below 85% of their normal body weight measured before the experiment. The mice were allowed to feed freely in their home cages. The experimental and housing protocols adhered to the Japanese National Regulations for Animal Welfare and were approved by the Animal Care and Use Committee of Keio University.

### Surgery

Mice were anesthetized with 1.0% to 2.5% isoflurane mixed with room air and placed in a stereotactic frame (942WOAE, David Kopf Instruments, Tujunga, CA, USA). A head post (H.E. Parmer Company, Nashville, TN, USA) was fixed at the surface of the skull, aligning the midline using dental cement (Product #56849, 3M Company, Saint Paul, MN, USA) to head-fix the mice during the experiment. The mice were group-housed (four mice per cage) before the experiments, and a recovery time of two weeks was scheduled between the surgery and experiment commencement.

### Procedure

Mice were habituated to a head-fixed experimental setup (Figure 1A; Kaneko et al., 2022; Toda et al., 2017; Yamamoto et al., 2022) the day before the experiment commenced. During habituation, mice were head-fixed in the apparatus with dim light and randomly presented with 10% sucrose solution through a drinking steel spout and a pure tone of 6000 Hz at 80 dB from a set of two speakers placed 30 cm in front of the platform. We conducted habituation to rewards and the auditory stimulus separately. The number of reward presentations during the habituation phase was not strictly defined; we determined that the mice were habituated by confirming that they consumed the reward stably from the spout by visibly checking the video. Auditory stimuli during the habituation phase were presented 120 times. Mice were head-fixed on a tunnel-like, covered platform by clamping a surgically implemented head plate on both sides (i.e., left and right from the anteroposterior axis of the skull). The clamps were placed on a slide bar next to the platform and adjusted to an appropriate height for each mouse. The platform floor was covered with a copper mesh sheet, and a touch sensor was connected to the sheet and steel spout.

**Figure 1.**
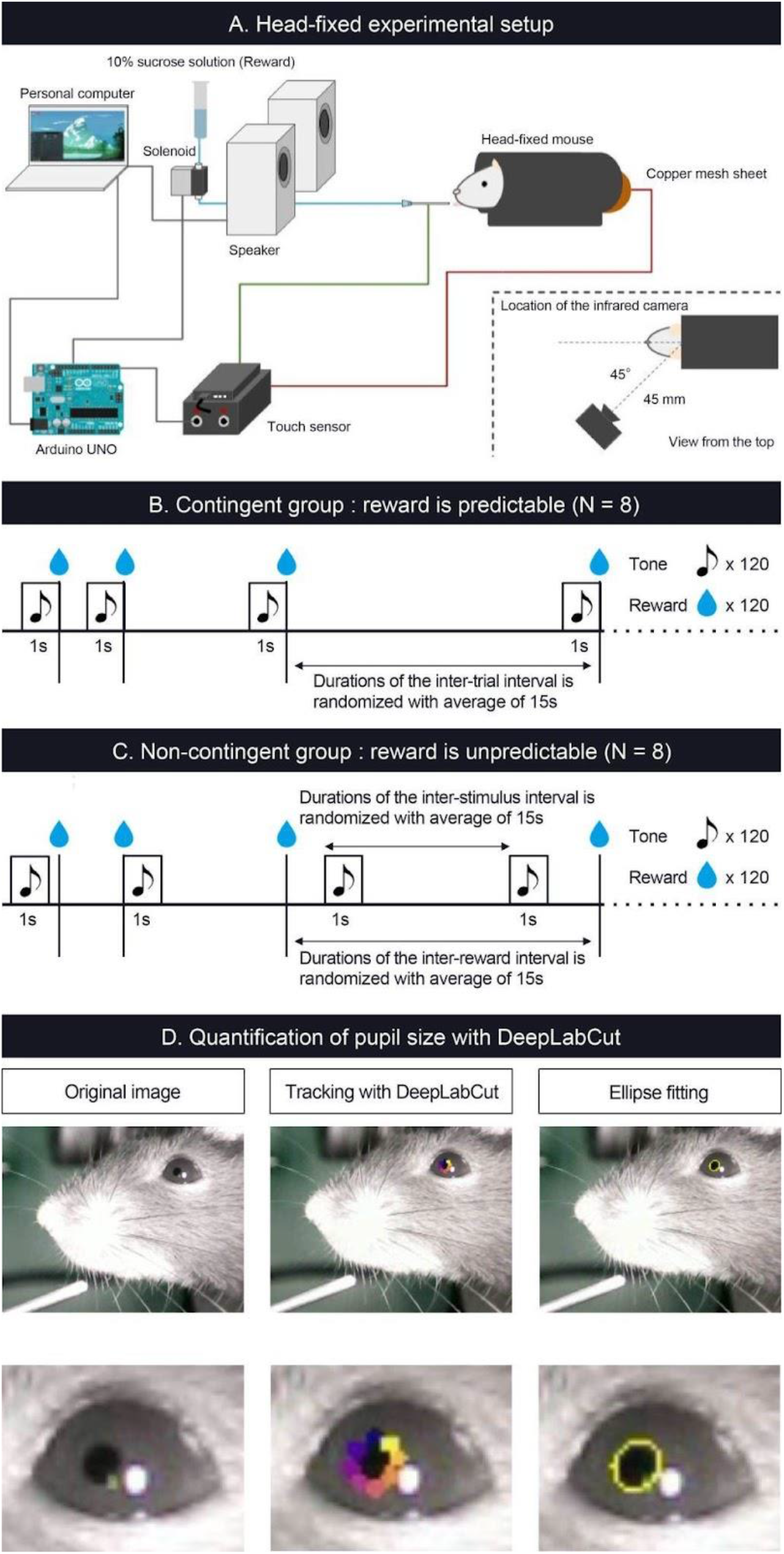
Schematic representation of head-fixed apparatus, Pavlovian delay conditioning task, and pupillometry. (A) Schematic representation of the head-fixed experimental apparatus and the custom-made experimental control system. (B) Contingent group. In this group, the 1 s auditory stimulus (6000 Hz tone) was followed by a reward delivery of a 4 μL drop of 10% of sucrose solution, and the auditory stimulus signaled the upcoming reward. (C) Non-contingent group. In this group, the auditory stimulus and the reward were presented independently and semi-randomly to prevent the development of the reward-predictive value of the auditory stimulus. (D) The left panel shows an image of a mouse’s eye taken by the infrared camera in mice performing the Pavlovian delay conditioning task. The center panel shows an image with eight tracked points using DeepLabCut. The right panel shows an image of an ellipse fitted to the points and an example of the temporal change in pupil size.

After habituation, we conducted a Pavlovian delay conditioning task. Figure 1 (B and C) shows the experimental procedure. Mice were assigned to two experimental groups, contingent (Figure 1B) and non-contingent (Figure 1C), with eight mice in each group. In the contingent group, a pure tone of 6000 Hz at 80 dB was randomly presented for 1 s as the conditioned stimulus (CS), followed immediately by a 4μl drop of 10% sucrose solution (Figure 1B). The CS presentation interval was random, ranging from 10 s to 20 s, and the mean value was set to 15 s. In the non-contingent group, the CS and reward were independently presented (Figure 1C). The CS and reward presentation intervals were random, ranging from 10 s to 20 s. We showed the percentages of CS and US overlapping trials for all individuals and sessions in the non-contingent group (Supplementary Figure 1), and there are no large differences between individuals. One session comprised 120 reward presentations for both groups. The training lasted for eight days. The CS and reward presentation, response, and video recording were controlled using a custom-made program written in Python 3 (3.7.8). The experiment was conducted in a soundproof box with 75 dB of white noise in the laboratory to mask external sounds.

### Drug

Pharmacological manipulations were conducted after training the Pavlovian conditioning task to suppress the licking response in mice. Six blocks were conducted for all individuals, each lasting three days. On Day 1, all mice were intraperitoneally administered saline solution 15 min before the experiment commenced. On Day 2, 15 minutes before the experiment commenced, haloperidol (Serenace, Sumitomo Pharma) 0.1, 0.2, and 0.5 mg/kg was administered intraperitoneally. This has been reported to inhibit licking (Fowler and Mortell, 1992; Liao and Ko, 1995) and spontaneous movements (Arruda et al., 2008; Bernardi et al., 1981; Conceição and Frussa-Filho, 1996; Strömbom, 1977) dose-dependently. After Day 2, mice were allowed to drink water freely for one hour. On Day 3, mice were not allowed access to water at all, and the experiment was not conducted to avoid the residual effects of the drug. Therefore, we set approximately 48 hours to wash out the effects of haloperidol. All individuals received each concentration of haloperidol twice. The administration followed an ascending (0.1, 0.2, 0.5, 0.1, 0.2, 0.5) and descending (0.5, 0.2, 0.1, 0.5, 0.2, 0.1) order. Four mice experienced the ascending order, and four experienced the descending order in each group. Haloperidol was diluted in a saline solution. We administered haloperidol to mice via intraperitoneal injection with a 10 mL/kg dose. After the injection, mice were returned to their home cage until the start of the experiment.

### Pupillometry

To measure the pupil size of mice performing the Pavlovian delay conditioning task, we used an infrared camera (Iroiro1, Iroiro House) to capture a video of the mice’s heads during the task. The camera was placed at 45° from the midline of the mouse (anteroposterior axis) and 45 mm from the top of the head (Figure 1A). The room’s brightness was set to 15 lux using a luminaire device (VE-2253, Etsumi). The pupil size was extracted from videos. Figure 1D shows the flow of the pupil size analysis. DeepLabCut, a deep-learning tracking software (Mathis et al., 2018; Nath et al., 2019), was used to track the pupil edge at eight points. Using the “EllipseModel” provided by scikit-image (Van der Wal et al., 2014), an ellipse was fitted to the eight points obtained by tracking, and the estimated parameters (major and minor diameters) were used to calculate the area of the ellipse. This area was used as pupil size. After fitting ellipses to the tracked points and calculating pupil area, each session’s data for all subjects were independently transformed to the standard normal distribution using the “scale” function in R. We employed “resnet50” as a backbone network of a model and used default parameters. We annotated 8 points, top, top-right, right, bottom-right, bottom, bottom-left, left, and top-left, for each frame. The dataset contains 1650 frames (15 frames per video and 110 videos). We trained the model with 1030000 iterations, and the train and test errors were 0.92 and 0.95 pixels, respectively.

### Licking bout analysis

Animal responses occur as bouts, characterized by bursts of responses and pauses that separate each bout (Gilbert, 1958; Shull et al., 2001). Conditioned responses (CR) also occur as bouts (Harris, 2015; Kirkpatrick, 2002; Toda et al., 2017). Since the CR has such a temporal pattern, individual licking can be classified into two types: those that occur within bursts and during pauses. In previous studies, such a bout-and-pause pattern was described by the mixture distribution of two exponential distributions (Killeen et al., 2002): *P*(*IRT* = τ) = *qe*^−*b*τ^ + (1 − *q*)*e*^−*w*τ^. In the equation, *q* denotes the mixture ratio of the two types of responses, and *w* and *b* denote the speed of the responses within bouts and the length of the pauses, respectively. These parameters were free, and we fitted the equation to the empirical data to estimate the parameters q, w, and b using a custom-made script and Turing (Ge et al., 2018), a Bayesian inference software in Julia language. Under the estimated parameters, individual licking was classified based on the likelihood of whether it occurred within bursts or during pauses.

### Statistical analysis

We collected data from all subjects repeatedly through our experiments and had missing values caused by the failure of video recording, and we excluded those data from the analysis. Given that our data was repeated-measurement data, including missing values, assumptions employed in standard statistical analysis (i.e., that data is independently and identically distributed) were violated. To account for repeated-measurement data, we employed a linear-mixed model that can be used in the same way as standard analysis of variance (ANOVA). Important aspects of the linear-mixed model are fixed and random effects. Fixed effects are of primary interest to researchers. In our experiments, group (contingent or non-contingent) and pharmacological treatment (dose of haloperidol) are fixed effects. Fixed effects can be interpreted analogously to effects examined in standard statistical analysis such as ANOVA. Random effects comprise additional variability from other sources, such as repeated measures clustered within subjects. By considering random effects, we can account for the effects of repeated measurement and assess the fixed effect more precisely (Singmann and Kellen, 2019). In statistical methodology, the maximal random effect structure is usually recommended to be modeled directly to reduce the Type I error when examining fixed effects (Barr et al., 2013). Thus, we included factors measured repeatedly within subjects (i.e., time windows, sessions, and pharmacological treatments) as random slopes. However, complex models have the risk of failing to converge and of overfitting (Bates et al., 2015). In such cases, several solutions have been proposed, and we have modified our implementation of the model in cases where we were facing convergence or overfitting problems (Brauer and Curtin, 2018). Therefore, we implemented a maximal random effect structure at first. When the model failed to converge or was overfitted to the data, we removed the random effect with the smallest variance. Finally, we employed the model in which the model converged without overfitting problems. The specific model we employed will be presented in the Results section. We fitted a linear mixed model to our data using R (4.1) and the *lme4* package (Bates et al., 2014). We also used the *emmeans* package (Lenth et al., 2018) to examine simple effects and employed Tukey’s method to adjust p-values for multiple comparisons.

## Results

In the contingent group, the stimulus signaled reward delivery. Licking and pupil responses increased after the auditory stimulus presentation and the sucrose solution delivery (“Contingent”; Figure 2B and 2C). In the non-contingent group, the auditory stimulus did not signal reward delivery. Licking and pupil responses did not change after the auditory stimulus presentation but increased after the sucrose solution delivery (“Non-contingent”; in Figure 2B and 2C). We set three periods for analysis of licking and pupil size, 1 s before the presentation of the auditory stimulus (Pre-CS period; shown in green the shade in Figure 2B), 1 s during the presentation of the auditory stimulus (CS period; shown in the red shade in Figure 2B), and 1 s after the reward presentation (US period; shown in the blue shade in Figure 2B). We computed linear-mixed models to examine the effects of the respective procedure on the amount of licking and pupil size at each time window. In the respective model, we included the time window (levels: Pre, CS, and US), group (levels: contingent and non-contingent), and their interaction as fixed effects, as well as random intercepts and random slopes for the variable time window on subject level. Linear-mixed for the amount of licking revealed significant interaction between the time window and the procedure (Left panel of Figure 2D; *F* (2, 13.999) = 105.8967, *p* <.0001). Subsequently, multiple comparisons revealed significant differences in the amount of licking between each time window in the contingent group (Contingent in the left panel of Figure 2D; Pre vs. CS: *t* (14) = −20.071, *p* <.0001; Pre vs. US: *t* (136) = −17.000, *p* <.0001; CS vs. US: *t* (136) = −4.033, *p* =.0033) and between Pre and US, US and CS in the non-contingent group (Non-contingent in the left panel of Figure 2D; Pre vs. US: *t* (14) = −15.275, *p* <.0001; CS vs. US: *t* (14) = −13.521, *p* <.0001), suggesting that the amount of licking increased after the auditory stimulus and reward presentation in the contingent group, but only after reward presentation in the non-contingent group. We also analyzed pupil size using an equivalent model, and the respective analysis revealed a significant main effect of the variable time window (Right panel of Figure 2D; *F* (2, 14.023) = 5.4407, *p* =.0178). Subsequently, we examined simple effects of the time window, which revealed significant differences in pupil size between Pre and US, CS and US in the contingent group (Right panel of Figure 2D; Pre vs. CS: *t* (13.8) = −2.563, *p* =.0557; Pre vs. US: *t* (14) = −3.346, *p* =.0125; CS vs. US: *t* (13.9) = −3.061, *p* =.0217), suggesting that pupil size increased after the reward presentation in the contingent group.

**Figure 2.**
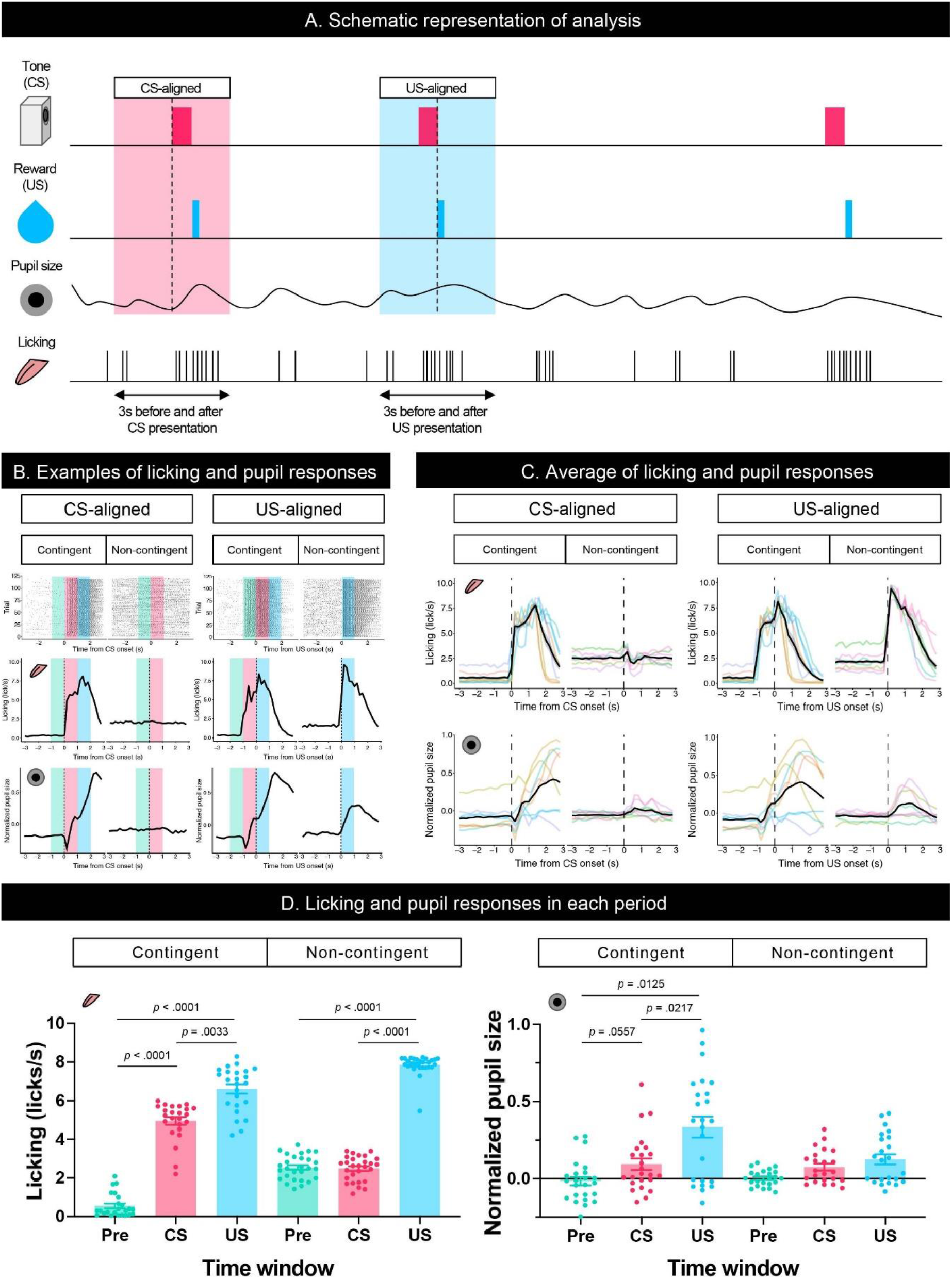
Results of the Pavlovian conditioning training. (A) Schematic representation of analyzed time windows. The presentation of CS or US was set as 0, and the 3 s before and after the presentation were used for analysis. (B). An example of licking responses and pupil response during each group’s Pavlovian delay conditioning task. Raster plot (top), temporal change in licking and pupil size (middle and low) of a representative individual from each of the contingent and non-contingent groups (*N* = 1 for each group, 120 trials each) 1 s before the auditory stimulus presentation (Pre), during the auditory stimulus presentation (CS), and immediately after the reward presentation (US) are shown in green, red, and blue, respectively. (C) Mean temporal changes in licking frequency and pupil size before and after CS and US presentations. Solid black lines indicate means; gray-covered areas indicate the standard error of the mean (*N* = 8 for each group, 3 sessions each). Thin, colored lines indicate individual data. (D) Licking frequency (left) and pupil size (right) at 1 s before, during, and immediately after CS presentations (*N* = 8 for each group, 3 sessions each). Each time window corresponds to the area covered by green, red, and blue in (A).

To examine the effect of licking responses on pupil size, we analyzed temporal changes in licking and pupil responses around the onset of the licking bout (Figure 3A). When we aligned the licking and pupil responses with the licking bout onset, both groups’ licking responses and pupil size increased with the bout onset (Figure 3B–D). Licking responses recorded a phasic increase at the bout onset, and the pupil size increased slightly after the bout onset. The pupil size slightly decreased before the bout onset and increased after the bout onset. These results indicate that pupil size increased after the initiation of licking responses. We examined the amount of licking and pupil size before and after bout onsets using a linear mixed model, which had the time window (Pre and Post) and group (contingent and non-contingent) as a fixed effect and random intercepts and random slopes of time window on subject level. The linear-mixed model revealed significant effects of the time window and group on the amount of licking (Left panel of Figure 3D; Pre vs. Post, *F* (1, 13.481) = 38.358, *p* < .0001; Contingent vs. Non-contingent, *F* (1, 13.948) = 5.5768, *p* = .0333), suggesting that the amount of licking increased after bout onset. The linear-mixed analysis also revealed a significant interaction of time window and group (Right panel of Figure 3D; *F* (1, 14.226) = 14.226, *p* = .0087) regarding pupil size. We examined the simple effects of time window on pupil size (Right panel of Figure 3D; Pre vs. Post, *t* (13.7) = −6.223, *p* < .0001 in the contingent group), suggesting that pupil size increased after bout onsets in the contingent group. We provide the results of fitting the mixture distribution of two exponential distributions in Supplementary Figure 2.

**Figure 3.**
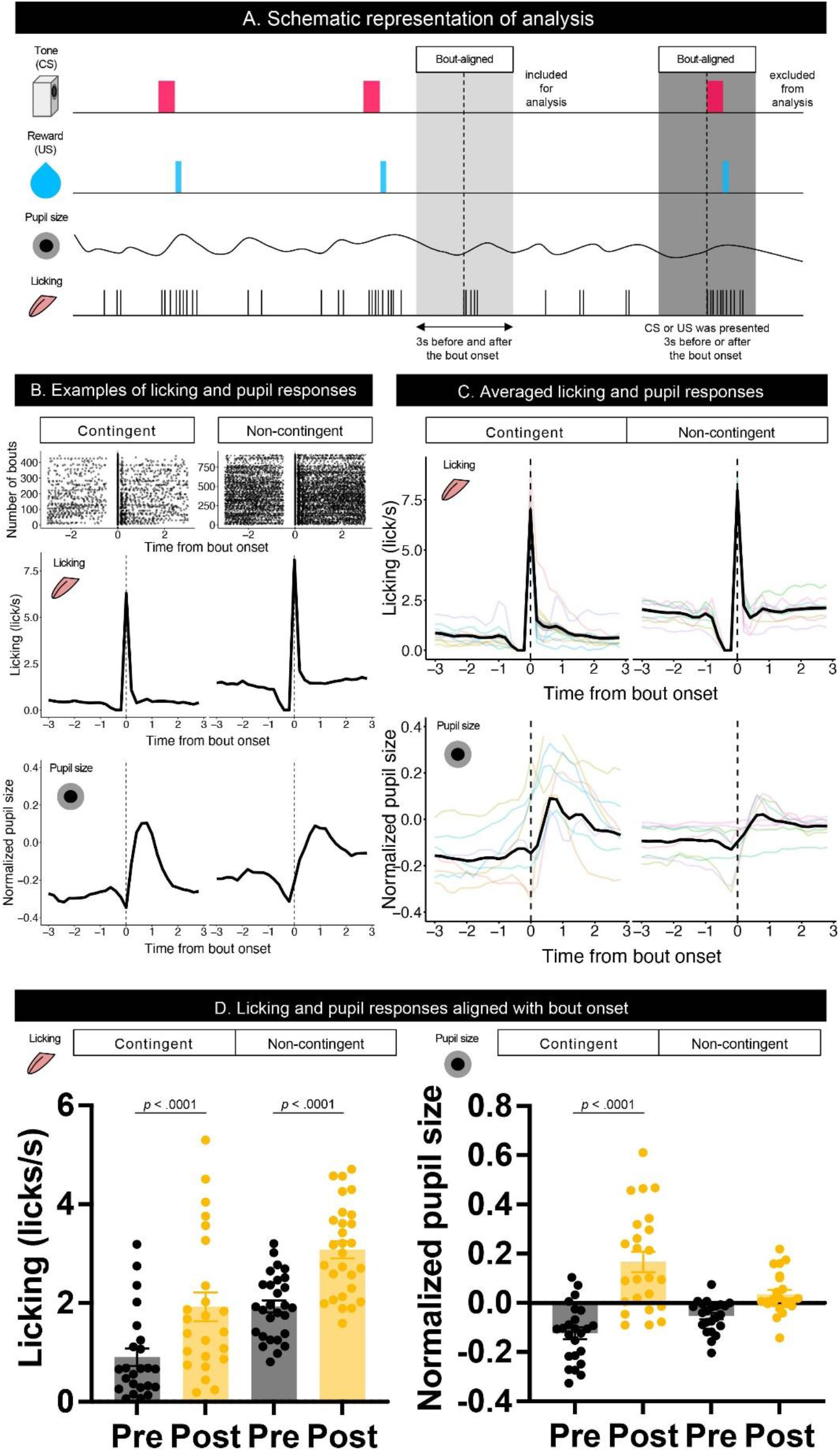
Temporal changes in licking responses and pupil size aligned with onsets of bouts. (A) Schematic representation of the response bout analysis. Initiation of the bout was set as 0, and the 3 s before and after bout initiation was used for the analysis. (B) Examples of raster plots before and after the start of the licking bout (top) and temporal changes in licking responses and pupil size (middle and low) in contingent and non-contingent groups (*N* = 1 for each group). (C) Average temporal changes in licking and pupil size in contingent and non-contingent groups (*N* = 8 for each group, 3 sessions each). (D) Mean amount of licking (left) and pupil size (right) at 3 s before and after the bout initiation (*N* = 8 for each group, 3 sessions each). In both (B) and (C), data, including CS and US presentations within 3 s before and after the initiation of the bout, were excluded. Individual data are shown as colored lines.

To further investigate whether the increase in pupil size resulted solely from licking responses or reward prediction independently of licking responses, we attempted to suppress licking responses in the same task. We thus intraperitoneally injected haloperidol, a dopamine D2 receptor antagonist known to suppress licking responses and locomotor activity (Fowler and Mortell, 1992; Liao and Ko, 1995; Strömbom, 1977; Bernardi et al., 1981; Conceição and Frussa-Filho, 1996; Arruda et al., 2008). After saline administration, licking responses and pupil size increased after the auditory stimulus presentation in the contingent group (Contingent in CS-aligned in Figure 4A and 4B) but remained unchanged in the non-contingent group (Non-contingent in CS-aligned in Figure 4A and 4B). We observed an increase in the licking frequency and pupil size at the reward delivery in both the contingent and non-contingent groups (US-aligned in Figure 4A and 4C). Systemic administration of haloperidol suppressed licking responses and pupil size in both the contingent and non-contingent groups (Figure 4A-D). In particular, the increase in pupil size after the reward delivery was suppressed in the contingent group (Contingent in US-aligned in Figure 4A and the solid line in Figure 4C). We examined the effects of group, time window, and dose of haloperidol on the amount of licking and pupil size using linear-mixed modeling where the time window (Pre, CS, and US), group (contingent and non-contingent), dose (saline, 0.0, 0.1, 0.2, and 0.5 mg/kg) and their interactions were considered fixed effects, while subject-level random intercepts and random sloped for the variables time window and dose were included. The linear-mixed model revealed significant interactions between all variables (Left panel of Figure 4D; *F* (6, 479.34) = 11.512, *p* < .0001), and we examined simple effects of the time window. In the contingent group, the amount of licking differed between Pre and CS, and Pre and US in all dose conditions (Upper left panel of Figure 4D; Saline: Pre vs. CS, *t* (23.7) = −19.044, *p* < .0001; Pre vs. US, *t* (16.0) = −12.864, *p* < .0001; 0.1 mg/kg: Pre vs. CS, *t* (83.2) = −9.364, *p* < .0001; Pre vs. US, *t* (25.2) = −6.877, *p* < .0001; 0.2 mg/kg Pre vs. CS, *t* (83.2) = −7.953, *p* < .0001; Pre vs. US, *t* (25.2) = −5.127, *p* = .0001; 0.5 mg/kg: Pre vs. CS, *t* (83.2) = −3.324, *p* = .0037; Pre vs. US, *t* (25.2) = −3.036, *p* = .0147), suggesting that the amount of licking increased after auditory stimulus presentations. In the non-contingent group, the amount of licking differed between Pre and US, and CS and US in all conditions (Bottom-left panel in Figure 4D; Saline: Pre vs. US, *t* (16.0) = −11.311, *p* < .0001; CS vs. US, *t* (16.2) = −11.729, *p* < .0001; 0.1 mg/kg: Pre vs. US, *t* (25.2) = −8.085, *p* < .0001; CS vs. US, *t* (26.4) = −8.038, *p* < .0001; 0.2 mg/kg: Pre vs. US, *t* (26.3) = −5.833, *p* < .0001; CS vs. US, *t* (27.7) = −5.603, *p* < 0.0001; 0.5 mg/kg: Pre vs. US, *t* (25.2) = −4.444, *p* = .0004; CS vs. US, *t* (26.4) = −4.413, *p* = .0004), suggesting that the amount of licking increased after reward presentations. We also analyzed pupil size with linear-mixed analysis, revealing a significant interaction between all variables (Right panel of Figure 4D; *F* (6, 479.52) = 3.009, *p* = .0068). We examined the simple effects of the time window and found significant differences between all the time windows in the contingent group (Upper right panel of Figure 4D; Saline: Pre vs. CS, *t* (20.7) = −5.450, *p* = .0001; Pre vs. US, *t* (17.6) = −8.389, *p* < .0001; 0.1 mg/kg: Pre vs. CS, *t* (59.0) = −3.452, *p* = .0029; Pre vs. US, *t* (36.0) = –7.569, *p* < .0001; 0.2 mg/kg Pre vs. CS, *t* (59.0) = −3.000, *p* = .0109; Pre vs. US, *t* (36.0) = −7.817, *p* < .0001; 0.5 mg/kg: Pre vs. CS, *t* (59.0) = −4.344, *p* = .0002; Pre vs. US, *t* (36.0) = 9.585, *p* < .0001), and Pre and CS in the non-contingent group (Bottom-right panel of Figure 4D; 0.1 mg/kg: Pre vs. CS, *t* (59.0) = −3.069, *p* = .0090; 0.2 mg/kg Pre vs. CS, *t* (63.8) = −2.579, *p* = .0324; 0.5 mg/kg: Pre vs. CS, *t* (59.0) = −2.496, *p* = .0402). The increase in licking responses and pupil size after the auditory stimulus presentation was examined by calculating the difference between the mean values of licking responses and pupil size for 3 s before and after the auditory stimulus presentation. We performed a linear-mixed analysis to the change in the amount of licking and pupil size. We assigned the group (contingent and non-contingent) and dose (saline, 0.1–0.5 mg/kg) to fixed effects and subject to random effect in licking analysis. Linear-mixed analysis revealed a significant interaction between the group and the dose. Subsequently, we examined the simple effect of dose and found that injection of haloperidol decreased the amount of licking in a dose-dependent manner in the contingent group (Upper left panel of Figure 4E; Saline vs. 0.1 mg/kg, *t* (169) = 7.741, *p* < .0001; Saline vs. 0.2 mg/kg, *t* (169) = 10.704, *p* < .0001; Saline vs. 0.5 mg/kg, *t* (169) = 16.205, *p* < .0001; 0.1 mg/kg vs. 0.5 mg/kg, *t* (169) = 6.910, *p* < .0001; 0.2 mg/kg vs. 0.5 mg/kg, *t* (169) = 4.492, *p* = .0001). We also analyzed pupil size using a linear mixed model, where group (contingent and non-contingent), dose (saline, 0.1–0.5 mg/kg), and the change in the amount of licking were considered fixed effects to assess pupil increase keeping out the effect of the increase of licks, and subject to the random intercept. Linear-mixed analysis revealed a significant effect of group, dose, and increase of licks (Upper right panel of Figure 4E; Group, *F* (1, 23.195) = 5.4185, *p* = .029; Dose, *F* (3, 173.005) = 3.1752, *p* = .0256; Licking, *F* (1, 184.886) = 4.7037, *p* = .0314), suggesting that pupil size showed stronger increases in the contingent group than the non-contingent group, even after ruling out effects of licks. We also examined the increase in licking responses and pupil size after examining the reward presentation by calculating the difference between the mean values of licking responses and pupil size for 3 s before and after the reward presentation. We performed linear-mixed analysis considering changes in the amount of licking and pupil size. We considered the variables group (contingent and non-contingent) and dose (saline, 0.1–0.5 mg/kg) as fixed effects and included subject-level random effects. Linear-mixed analysis revealed a significant interaction between the group and the dose. Subsequently, we examined the simple effects of dose and found that injection of haloperidol decreased the amount of licking in a dose-dependent manner (Contingent in the bottom-left panel of Figure 4E; Saline vs. 0.2 mg/kg, *t* (169) = 3.878, *p* = .0009; Saline vs. 0.5 mg/kg, *t* (169) = 4.220, *p* = .0002; Non-contingent in the bottom-left panel of Figure 4E; Saline vs. 0.1 mg/kg, *t* (169) = 3.052, *p* = .0139; Saline vs. 0.2 mg/kg, *t* (169) = 6.093, *p* < .0001; Saline vs. 0.5 mg/kg, *t* (169) = 8.476, *p* < .0001; 0.1 vs. 0.5 mg/kg, *t* (169) = 4.429, *p* = .0001). We also analyzed pupil size using a linear mixed model, where group (contingent and non-contingent), dose (saline, 0.1–0.5 mg/kg), and the change in the amount of licking were considered fixed effects to assess pupil increase keeping out the effect of the increase of licks, and subject to the random intercept. Linear-mixed analysis revealed a significant effect of group (Bottom-right panel of Figure 4E; Group, *F* (1, 18.395) = 8.683, *p* = .0085), suggesting that pupil size significantly more strongly increased in the contingent group than the non-contingent group.

**Figure 4.**
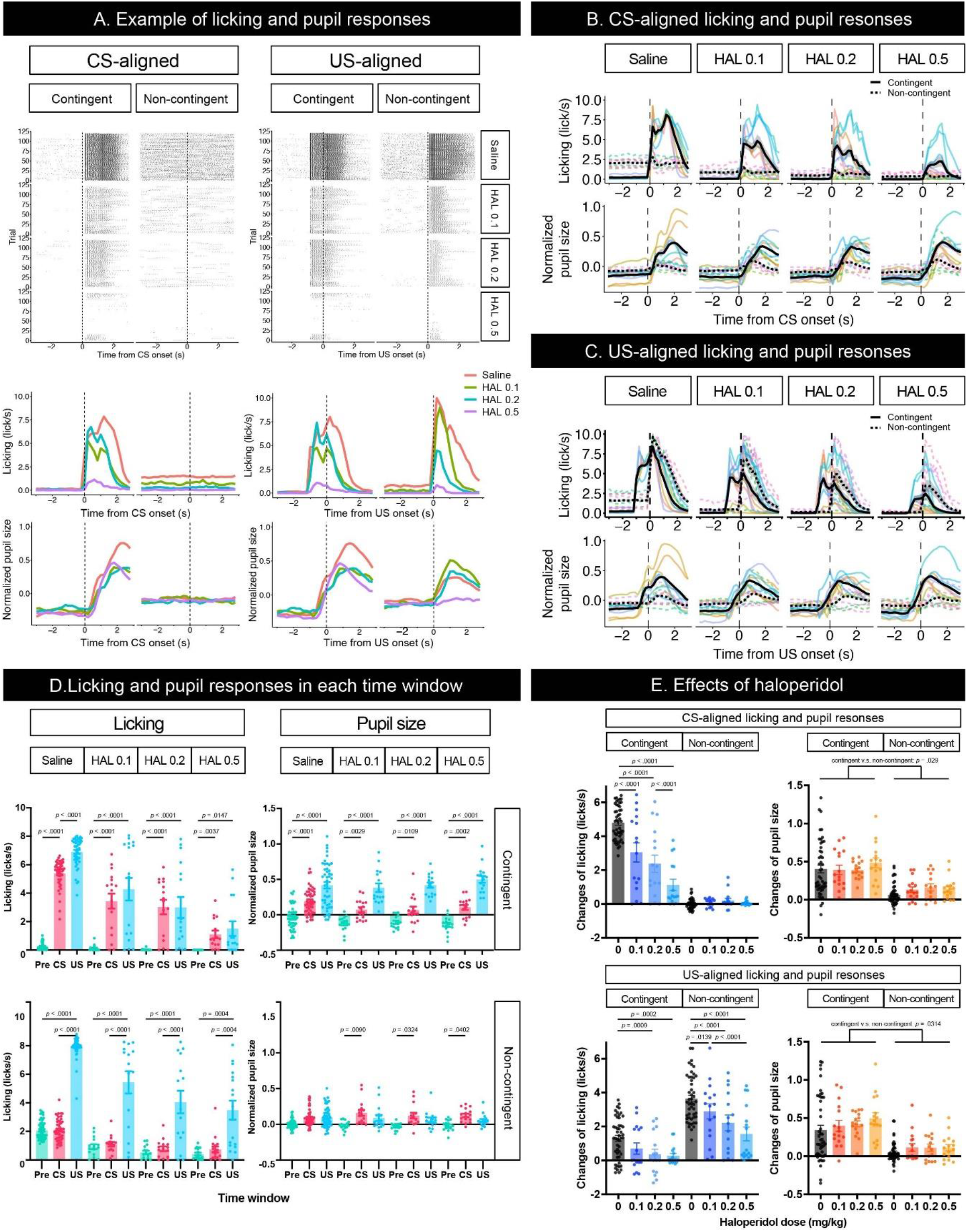
Effects of haloperidol injection on the licking and pupil responses after Pavlovian conditioning training. (A) Representative raster plot (top), temporal change in licking, and pupil responses (middle and low) of individuals in contingent and non-contingent groups. Periods of 1 s before the auditory stimulus presentation (Pre-CS), during the auditory stimulus presentation (CS), and after the reward presentation (US) are shown in green, red, and blue, respectively (*N* = 1 for each group, 120 trials each). (B) Mean temporal changes in licking and pupil size before and after CS presentations (*N* = 8, 6 sessions for saline condition and 2 sessions for all haloperidol conditions). The upper panel indicates licking responses. The horizontal axis indicates the time from the reward onset. The vertical axis indicates frequencies of licking responses. The lower panel indicates the data on pupil size. The horizontal axis indicates the time from the reward onset. The vertical axis indicates the normalized pupil size. (C) Mean temporal changes in licking and pupil responses before and after US presentations. (D) Licking responses at Pre-CS, CS, and US periods (*N* = 8, 6 sessions for saline condition and 2 sessions for all haloperidol conditions). The pupil size at Pre-CS, CS, and US periods. (E) Difference between the mean values of licking responses and the normalized pupil size during a 3 s before and after CS presentation (upper panel) and reward presentation (bottom panel). HAL indicates haloperidol.

## Discussion

This study explored the dynamics of the licking response and pupil size while mice performed a Pavlovian delay conditioning task to investigate the relationship between reward prediction and pupil size. The head-fixed experimental setup combined with deep-learning-based image analysis enabled us to reduce mice’s spontaneous locomotor activity and to track the precise dynamics of licking responses and pupil size of the behaving mice. By manipulating the predictability of the reward in the Pavlovian delay conditioning task, we demonstrated that the pupil size of mice was modulated by reward prediction, consumption of the reward, and body movements associated with reward processing. Additionally, we found that the pupil size was modulated by reward prediction even after dose-dependent disruption of body movements by intraperitoneal injection of haloperidol, a dopamine D2 receptor antagonist.

In Experiment 1, we trained head-fixed mice on the Pavlovian delay conditioning task while recording licking and pupil responses. In this task, we designed contingent and non-contingent conditions to manipulate the predictability of the delivery of the sucrose solution by the auditory stimulus. In the contingent group, the auditory stimulus signaled the sucrose solution delivery. The mice showed increased licking responses and pupil size after the auditory stimulus presentation, suggesting that they could predict the outcome in this group. In the non-contingent group, the auditory stimulus did not signal the reward delivery. Licking responses and the pupil size of mice remained unchanged by the auditory stimulus presentation, suggesting that they did not associate the auditory stimulus with the reward in this group. In addition, the behavioral results obtained from the non-contingent group demonstrated that the sensory stimulus itself did not affect changes in licking responses and pupil size. The frequencies of the auditory stimulus presentation and reward delivery were identical between the contingent and non-contingent groups, with the only difference being the predictability of the outcome following the auditory stimulus. This well-controlled rigid behavioral design allowed us to investigate the modulation of behavioral states induced by reward prediction with the same sensory signals.

Detailed bout analysis of licking responses revealed that pupil size increased after the licking bout initiation in both the contingent and non-contingent groups, suggesting that licking responses may modulate pupil size. Bout-aligned pupil size also showed a clear decrease before the increase in pupil size. Before the bout initiation, there was no licking response for approximately 0.5 s (Figure 3B and 3C). This result also confirms the close relationship between pupil size and licking responses. Many kinds of anticipatory behaviors occur when the stimulus signals a future outcome. Thus, whether changes in pupil size reflect signals related to reward prediction or are simply modulated by the motor-related signals accompanied by the predictive movement remains unclear.

To examine whether the changes in pupil size reflect the modulations by the prediction irrespective of motor-related signals, we examined the effects of intraperitoneal injection of haloperidol, a dopamine D2 receptor antagonist, on the dynamics of the pupil size of mice performing the Pavlovian delay conditioning task in Experiment 2. Intraperitoneal injection of haloperidol suppressed licking responses in a dose-dependent manner, supporting previous findings (Fowler and Mortell, 1992; Liao and Ko, 1995). Although haloperidol administration decreased pupil size, the effect was not as drastic as that of licking responses (Figure 4D and 4E). The highest dose of haloperidol injection almost completely disrupted licking responses; however, we still observed pupil dilation after the auditory stimulus presentation in the contingent group. This result implies that changes in pupil size reflect reward-predictive signals irrespective of movement-related modulations.

In our experiments, the pupil size and licking responses were larger in the non-contingent group, even if CSs were not presented compared to the contingent group (Figure 4C and 4D). The timing of the reward presentations was completely unpredictable in the non-contingent group, but the context predicted the possibility of reward presentations. Pupil size was not increased after unpredictable reward presentation in the non-contingent condition, suggesting that pupil size did not reflect reward prediction errors in our experimental context. In our experiments, subjects were trained extensively in a non-contingent context. This overtraining condition might have caused the subjects to have no prediction errors at reward presentation, even if the rewards were unpredicted in the non-contingent group. In further studies, investigating the effect of reward prediction errors or uncertainty on pupil size by presenting or omitting the reward will be important to understand the relationship between pupil size changes and reward prediction errors.

In this study, we explored the dynamics of pupil size of mice performing the Pavlovian delay conditioning task. We found that pupil dynamics reflected reward prediction signals, irrespective of modulations by body movements. Pupil size is modulated by autonomic nervous system activity. Sympathetic and parasympathetic activation lead to pupil expansion and contraction, respectively. The sympathetic control of the pupil is mediated by neuronal activity in the intermediolateral cell column (IML) of the cervical and thoracic regions of the spinal cord. Cholinergic neurons mediate the parasympathetic control in the Edinger-Westphal nucleus (EWN). Most locus coeruleus (LC) neurons are noradrenergic, and their direct projections to the IML stimulate sympathetic activation via noradrenergic α1 receptors. Direct projections to the EWN are thought to suppress the parasympathetic nervous system by acting in an inhibitory manner via α2 receptors (Joshi and Gold, 2020). Simultaneous measurements of LC neuronal activity and pupil size in monkeys and rats have been reported to correlate (Joshi et al., 2016; Liu et al., 2017). Therefore, pupil size measurement can be interpreted as an indirect measure of LC activity.

Considering that LC neuronal activity is highly correlated with pupil size (Joshi et al., 2016; Liu et al., 2017), our results that (1) the pupil was dilated to the auditory stimulus that predicted the reward, and (2) the pupil size was unchanged when the stimulus signaled no information about the reward, are consistent with existing findings from electrophysiological experiments of LC neurons (Aston-Jones et al., 1997, 2005; Bouret and Richmond, 2015; Bouret and Sara, 2004). LC neurons show a burst of activity when the stimuli that predict biologically important events, such as reward and aversive events, are presented (Aston-Jones ad Bloom, 1981; Aston-Jones et al., 1997, 2005; Bouret and Richmond, 2015; Bouret and Sara, 2004). LC neurons also show similar activities to dopaminergic neurons in the ventral tegmental area (VTA), such as increased phasic activity in response to unpredicted reward and decreased activity through repeated experience and transfer to a reward-predicting stimulus (Amo et al., 2022; Bouret and Sara, 2004; Schultz et al., 1997). However, the neuronal activities of LC neurons in the study of Bouret and Sara (2003) were examined with the reversal of the contingency between the stimulus and the outcome or the re-acquisition after the extinction in the Go/No-Go task. Although the phasic activities to unpredicted reward found in LC neurons (Bouret and Sara, 2004) may be slightly different from those neurons found in dopaminergic neurons found in VTA (Amo et al., 2022; Schultz et al., 1997), both phasic activities of LC and DA neurons are known to show phasic responses to unpredictable events. Moreover, LC neurons show phasic activity in response to a novel stimulus and decreased activity when the stimulus ceases to predict biologically important events (Berridge and Waterhouse, 2003; Vankov et al., 1995). LC activity might be related to the salience-related dopaminergic activity found in the midbrain (Matsumoto and Hikosaka, 2009). As shown in Figures 2B and 4B, pupil size dilated for the reward-predictive stimulus but not for the reward-non-predictive stimulus. Although the reward prediction error did not modulate pupil size, the dynamics of pupil size observed in our experiments could be partially interpreted as reflecting LC activity.

In the canonical view of the reward prediction error hypothesis, neuronal activities of dopamine neurons in the VTA are modulated by the reward prediction errors and this signal is considered as teaching signals (Bayer and Glimcher, 2005; Eshel et al., 2016; Hollerman and Schultz, 1998; Satoh et al., 2003; Schultz et al., 1997). Learning also involves several other components, such as modulation of motor outputs. Pupil size is considered to reflect internal states of organisms involving arousal and/or attention and is modulated by noradrenergic neurons in the LC. We found that haloperidol suppressed licking responses but not pupil size, suggesting that dopamine D2 receptors are not involved in the modulation of the reward prediction itself or attention/arousal modulated by the reward prediction. In contrast, the fact that licking responses are suppressed by haloperidol suggests that dopamine D2 receptors play a crucial role in the motor output based on the reward prediction. Theoretically, if output signals of the reward prediction error are modulated by the manipulation, the prediction itself as “associative strength” assumed in the associative learning theory or “value” assumed in the reinforcement learning theory would be also updated. In this sense, our results suggest that the output of dopaminergic signals from the midbrain to D2 receptors in the brain areas that receive dopaminergic projections might have an important role in modulating the motor output irrespective of updating the reward-predictive signals. This conclusion from our study supports recent findings that showed the neuronal activities of dopamine neurons in the midbrain encode information about movement kinematics (Barter et al., 2015; Hughes et al., 2020). However, we did not examine the effect of the other dopamine receptor, dopamine D1 receptor. In the future, examining the pharmacological manipulation of dopamine D1 receptors is an important step for better circuit-level understanding of the neuronal mechanism of the reward prediction and the reward prediction error.

Dopamine neurons in VTA show phasic activity to unexpected reward presentations, but phasic activity to the reward decreases as learning progresses, and the neurons show phasic activity to reward-predictive cues (Amo et al., 2022; Schultz et al., 1997). Thus, in our experiments, increases in pupil size may reflect reward prediction errors in the presentation of the auditory stimulus. However, increases in pupil size for reward delivery were small in the non-contingent group (Figure 2C, D and 4C, D). In addition, if the pupil size is modulated by the reward prediction errors, pupil dilation should occur after the reward presentation only in the non-contingent group. In our experiments, however, pupil dilation after reward presentations also occurred in the contingent group where reward presentations could be fully predictable by the auditory stimulus. These results suggest that the pupil size did not reflect reward prediction errors in our experiments. In addition, human studies indicated that the pupil size dilated for the reward-predictive cue in a delay conditioning task where the cue was presented 5 s before the reward (Pietrock et al., 2019). Taken together, increases in pupil size caused by the presentation of auditory stimuli in the contingent group could be interpreted as reward-predictive signals.

Considering the neurobiological mechanisms underlying the pupillary control system, the present findings of changes in pupil size being reflective of reward prediction signals invite us to reconsider the neuronal circuits computing reward prediction error signals. Cohen et al. (2012) reported that neuronal activities of GABAergic neurons in the rodent’s VTA reflect the prediction of upcoming reward values. These activities are considered the source of the prediction for computing reward prediction errors encoded in dopamine neurons in the VTA. In the study of Cohen et al. (2012), they recorded neuronal activity while the mice performed a Pavlovian trace conditioning task, in which each odor cue was associated with different upcoming outcomes, e.g., small and large amounts of liquid rewards and air puffs. GABAergic neurons in the VTA showed persistent ramping activity during the delay between the presentation of cues and reward. However, CR, such as licking the reward spout, occurred during the delay between the cue and the reward delivery. In such cases, it is difficult to assess whether the neuronal activity reflects the reward value or behavioral expression, for example, the motor activity involved in licking responses modulated by the reward value. In the present study, we attempted to overcome this problem by suppressing body movements with haloperidol and found that the changes in pupil size reflected reward prediction signals independent of licking movements. The integrative approach of behavioral analysis, image analysis, pupillometry, and pharmacological manipulations employed in the present study will pave the way for understanding the psychological and neurobiological mechanisms involved in the computation of reward prediction and reward prediction errors, which are essential features of learning and behavior.

We identified three limitations in this study: (1) the influence of body movements other than licking responses, (2) the pharmacological selectivity to haloperidol, and (3) the properties of CS and US, such as duration, magnitude, and timing, being determinants of learned responses. In appetitive Pavlovian conditioning, the presentation of the cue that predicts the outcome leads to the observation of approach behavior to the cue or to the location where the reward is presented (Boakes, 1977; Hearst and Jenkins, 1974). The locomotor activity also occurs in mice under a head-fixed situation and has been reported to affect pupil size (Cazettes et al., 2021). Intraperitoneal injection of haloperidol has been known to dose-dependently decrease spontaneous activities, including locomotor activities. Therefore, we hypothesized that the effect of locomotion on pupil size would be low in our experiments. However, we found that licking responses were not suppressed in all subjects. We cannot exclude this possibility because we could only measure licking responses and no other motor expressions in our head-fixed setup. Taken together, our experiments are limited mainly due to the potential effect of body movement on pupil size. Second, we used haloperidol to suppress mice’s body movements, but haloperidol might affect pupil size due to its non-selective nature. Haloperidol is a non-selective dopamine D2 antagonist that binds to D2-like receptors, including D3 and D4 receptors, and others, such as adrenergic α1 receptors. Adrenergic α1 receptors are involved in pupil dilation, and haloperidol has been reported to suppress pupil dilation produced by adrenaline administration in mice (Korczyn and Keren, 1980). Furthermore, electrical stimulation of the LC triggers the activity of dopamine cells in the midbrain via adrenergic α1 and dopamine release in the nucleus accumbens (Grenhoff et al., 1993; Park et al., 2017). Since pupil size is highly correlated with LC activity (Joshi et al., 2016; Liu et al., 2017), the injection of haloperidol may affect the activity of dopamine neurons in the midbrain and nucleus accumbens, which are modulated by LC activity. This suggests that haloperidol may consequently affect reward prediction and the calculation of reward prediction error. Furthermore, since it has been reported that body movements suppress the activity of the auditory cortex in mice (Nelson et al., 2013), it is possible that the injection of haloperidol suppresses body movement and consequently modulates pupillary responsiveness to the CS. The results of Experiment 2 may reflect this factor, where pupil dilation to CS was observed in the non-contingent group after haloperidol injection.

In this study, we used haloperidol, a non-selective Dopamine D2 antagonist. Although haloperidol does not increase pupil size, we might obtain cleaner results if a more selective antagonist is used. In future investigations, the use of selective dopamine D2 antagonists, such as eticlopride, and in combination with selective dopamine D1 antagonists, such as SCH-23390, may more definitely prevent modulations by the motor output and refine our understanding of the neurobiological mechanisms underlying the relationship between the pupil size and reward prediction.

Properties of CS and US, such as duration, intensity, and timing, affect learned responses (Fanselow, 1994; Holland, 1977; Solomon and Corbit, 1974; Timberlake, 1994). Taking such characteristics of conditioning, these factors may have affected the result of our study. In our experiments, the duration of the auditory stimulus was short, and the reward was followed by the auditory stimulus immediately. In such a situation, mice would gain their arousal immediately after the auditory stimulus presentation to consume the reward immediately. However, if the auditory stimulus duration was long, the CS presentation was not followed by the reward immediately, mice do not need to prepare to consume the reward immediately after the CS presentation, and this kind of difference in the task structure may lead to a different result. Although we did not use different types of US in our experiments, pupil size may show that CR differs depending on CS and US properties.

To verify that organisms predict future outcomes, behavioral evidence of preparatory or anticipatory responses is necessary. In general, anticipatory responses are accompanied by motor expressions; thus, dissociating whether the physiological changes related to reward prediction encode the signal of the prediction itself or are simply modulated by motor-related signals is difficult. Here, we successfully measured changes in pupil size in mice performing the Pavlovian delay conditioning task in the head-fixed situation using image processing. We revealed that dynamic changes in pupil size reflect reward-predictive signals. Pharmacological intervention experiments using haloperidol demonstrated that pupil size increased even when licking responses were suppressed, supporting that the changes in pupil size reflect reward prediction. Considering the brain circuits involved in controlling pupil size, the predictive feature of pupil size suggests that reward prediction is encoded in regions other than those reported by Cohen et al. (2012) and Tian et al. (2016). These results pave the way for our understanding of reward prediction signals in the brain by neutralizing the factor of motor expression and suggest a different hypothesis for the neuronal circuits of predictive learning. Future studies are expected to identify the neuronal circuit that computes the reward prediction and reward prediction errors by eliminating the modulation of motor expressions.

## Data availability statement

Data supporting the findings of this study are available from the corresponding author upon reasonable request. The original codes written for the analysis are available from the corresponding author upon reasonable request.

## Ethics statement

The experimental and housing protocols adhered to the Japanese National Regulations for Animal Welfare and were approved by the Animal Care and Use Committee of Keio University.

## Author contributions

YK and KT designed the experiments. YK conducted stereotaxic surgery and pharmacological manipulations for mice and collected all the data from the head-fixed Pavlovian conditioning experiment with the help of KT. YK and KT analyzed the data and created all figures. YK and KT wrote the manuscript.

## Funding

This research was supported by JSPS KAKENHI 18KK0070(KT), 19H05316 (KT), 19K03385 (KT), 19H01769 (KT), 20J21568 (KY), 22H01105 (KT), Keio Academic Development Fund (KT), and Keio Gijuku Fukuzawa Memorial Fund for the Advancement of Education and Research (KT).

## Conflict of interest

The authors declare that they have no conflict of interest.

## Acknowledgments

We thank Kohei Yamamoto, Saya Yatagai, Yusuke Ujihara, Daiki Nasukawa, Yasuyuki Niki, Haruka Hirakata, Shohei Kaneko, Ryuto Tamura, Lingcheng Kong, and Kazuko Hayashi for their assistance with animal care and Youcef Bouchekioua, Akihiro Funamizu, and Takaaki Ozawa for valuable discussions. We also thank the anonymous reviewers for their constructive comments on the manuscript.

## Supplementary Materials

**Supplementary Figure 1.**
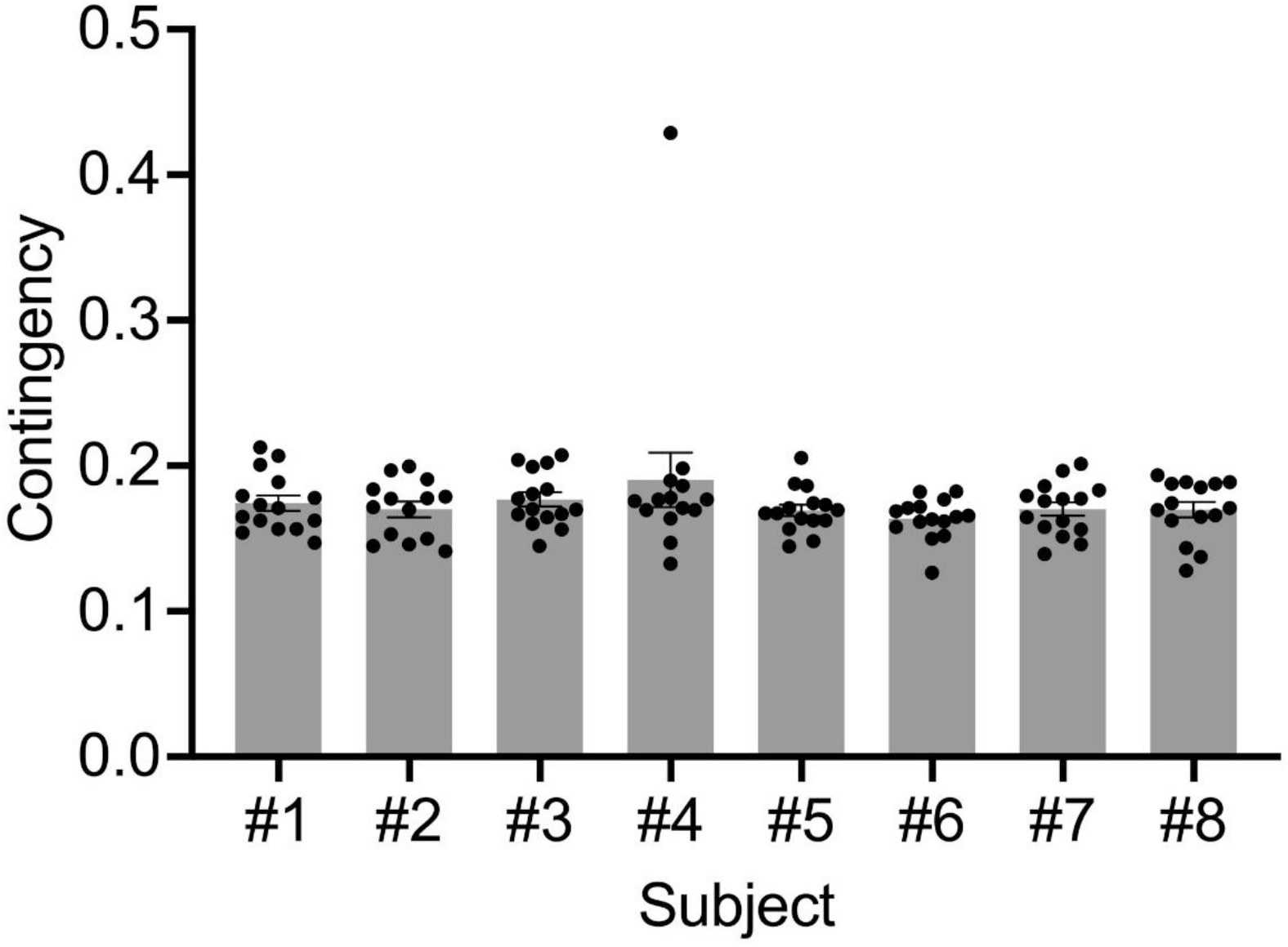
Empirical contingency in the non-contingent group. We calculated the percentage of CS and US overlapping trials for all individuals and sessions in the non-contingent group. The definition of overlap was determined by whether reward presentations were presented during auditory stimulus presentations.

The log survivor plot is a method to visualize the bout-and-pause patterns, and when responses have bout-and-pause patterns, the plot shows the broken-stick curve. The left side line denotes the within-bout inter-licking intervals, and the right-side line denotes the bout initiation intervals. The intercept of the right-side line denotes the bout length, the amount of licking contained in one bout. Log survivor plots of empirical and simulated data showed broken-stick curves, suggesting licks have bout-and-pause patterns. As shown in the figure, the bend points were approximately 0.1–1.0s, suggesting that the boundaries separating the within-bout licking from the bout initiation licking were in the range and corresponded to Figure 3B and C.

We found that the log survivor plot in the contingent group showed a clear bend point in training and saline conditions; in contrast, the plot did not show a clear bend point and showed a gradual curve in the non-contingent group. As the dose of haloperidol increased, the bend point became clearer. In the contingent group, the slope of the right lines became gradual as the dose of haloperidol increased. Taken together, mice did not show spontaneous licking during the inter-reward intervals.

**Supplementary Figure 2.**
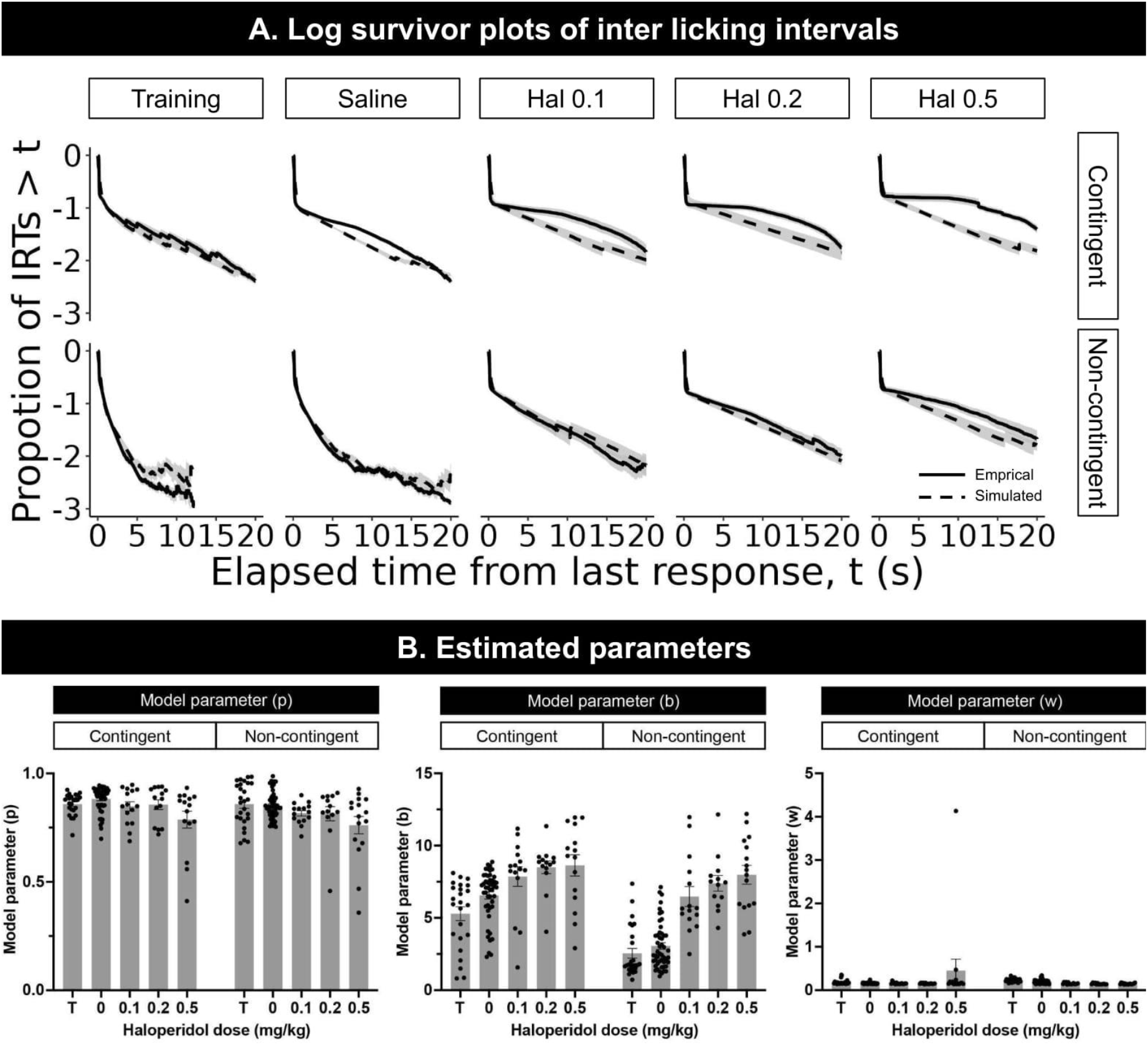
Fitting results of the mixture exponential distribution. (A) Solid lines show log survivor plots of empirical inter-licking intervals, and dashed lines show log survivor plots of simulated data using fitted parameters. Model fitting was performed independently for individual and session data, and we generated random numbers from the distribution of the estimated parameters. Each line denotes the average over subjects and sessions, and gray shades denote standard errors. (B) Average value and range of estimated parameters, *w*, *b*, and *p*, in each group and dose condition. T denotes the data in the last three sessions of training.

We compared the dynamics of licking and pupil size before/after the auditory stimulus presentation between the first and last training sessions. However, we failed to record several data in the first session, showing only 4 and 3 subjects for contingent and non-contingent groups. In the first session, the amount of licking slightly increased after the auditory stimulus in the contingent group but not in the non-contingent group. In the last session, the increase in the amount of licking became larger in the contingent group. In the non-contingent group, the amount of licking did not increase after the auditory stimulus presentation, but the baseline was larger in the last session compared to the first session. In both groups, mice showed pupil dilation to the auditory stimulus in the first session, but in the last session, it decreased in the non-contingent group but not in the contingent group.

The amount of licking was acquired by Pavlovian conditioning, but the pupil size showed an increase in the very first session. The pupil size is highly correlated with LC activity, and LC shows the phasic activity to a novel stimulus. It also shows the activity when the environmental rule, such as stimulus-reward contingency, changes. The increase in pupil size in the first session may reflect the novelty of stimulus or change in the environmental rule, such as the transition from habituation to contingent or non-contingent procedure. At least, pupil size does not reflect only reward prediction but also novelty, uncertainty, and environmental changes, so that a well-known learning curve may not be drawn.

**Supplementary Figure 3.**
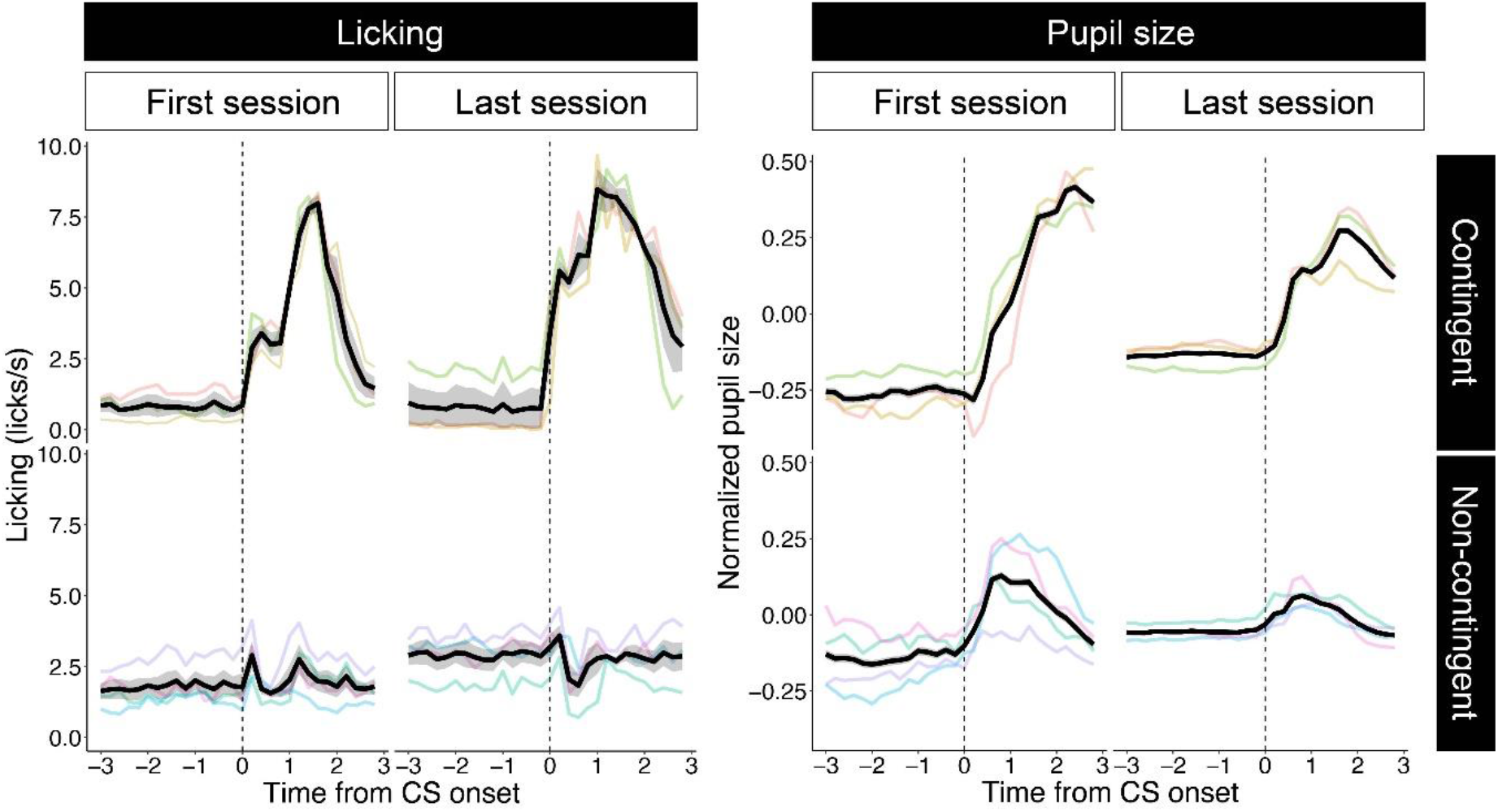
Comparison of dynamics of licking and pupil size between early and last session in training.

We analyzed the relationship between the amount of licking and pupil size in each trial. Although licking increased pupil size, as shown in Figure 3, we could not find any relationship between the amount of licking and pupil size. The large temporal variance in pupil size may mask the relationship between licking and pupil size in this time-scale analysis.

**Supplementary Figure 4.**
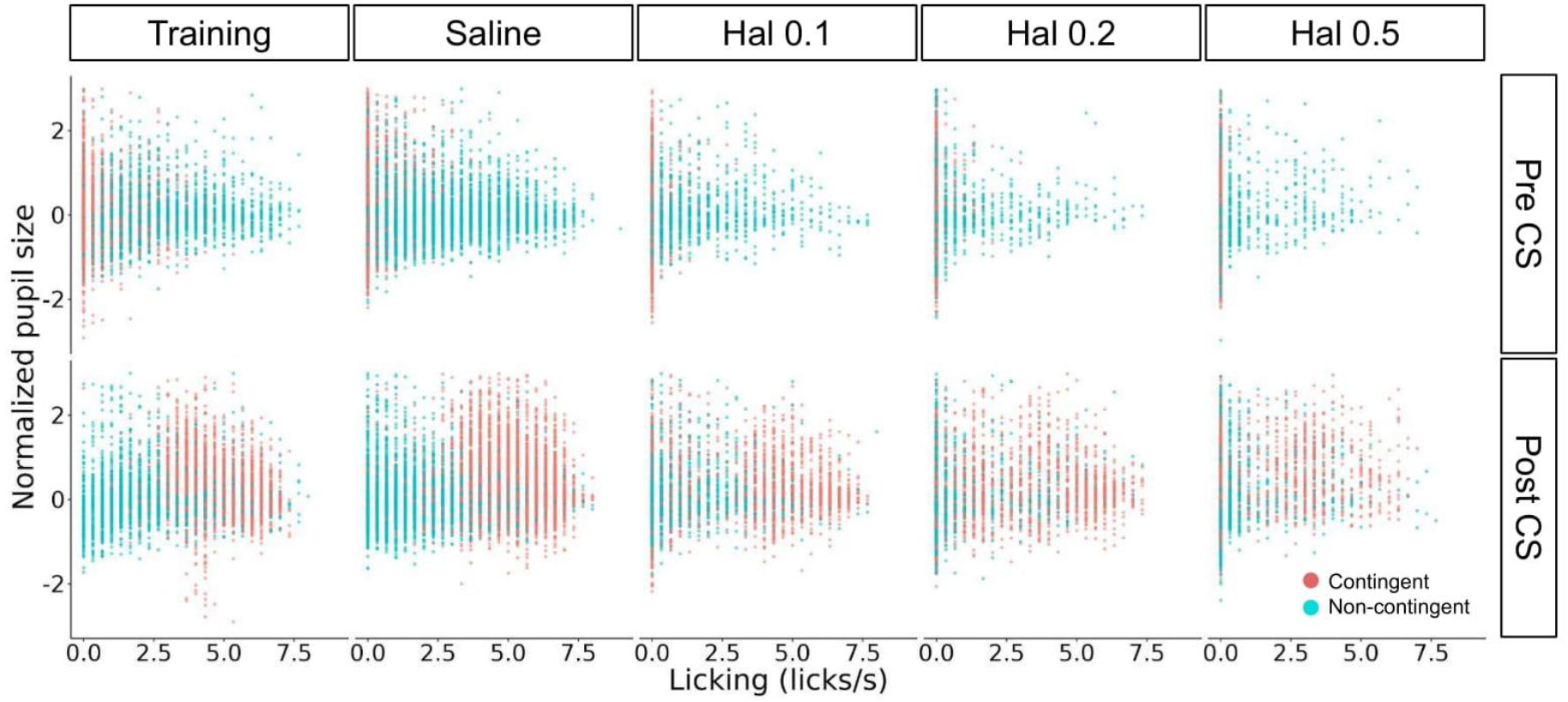
Scatter plots of the amount of licking and pupil size. The upper panels show relationships between the amount of licking and pupil size at 3 s before the auditory stimulus presentation. The bottom panels show those of 3 s after the auditory stimulus presentation. Blue and red points denote the non-contingent and contingent groups, respectively. We analyzed the saline condition separately for previous haloperidol dose conditions to examine whether the effects of haloperidol were washed-out. We found no difference between the previous dose in the amount of licking and pupil size at any time window, suggesting that the effects of haloperidol had been washed-out until the saline condition.

**Supplementary Figure 5.**
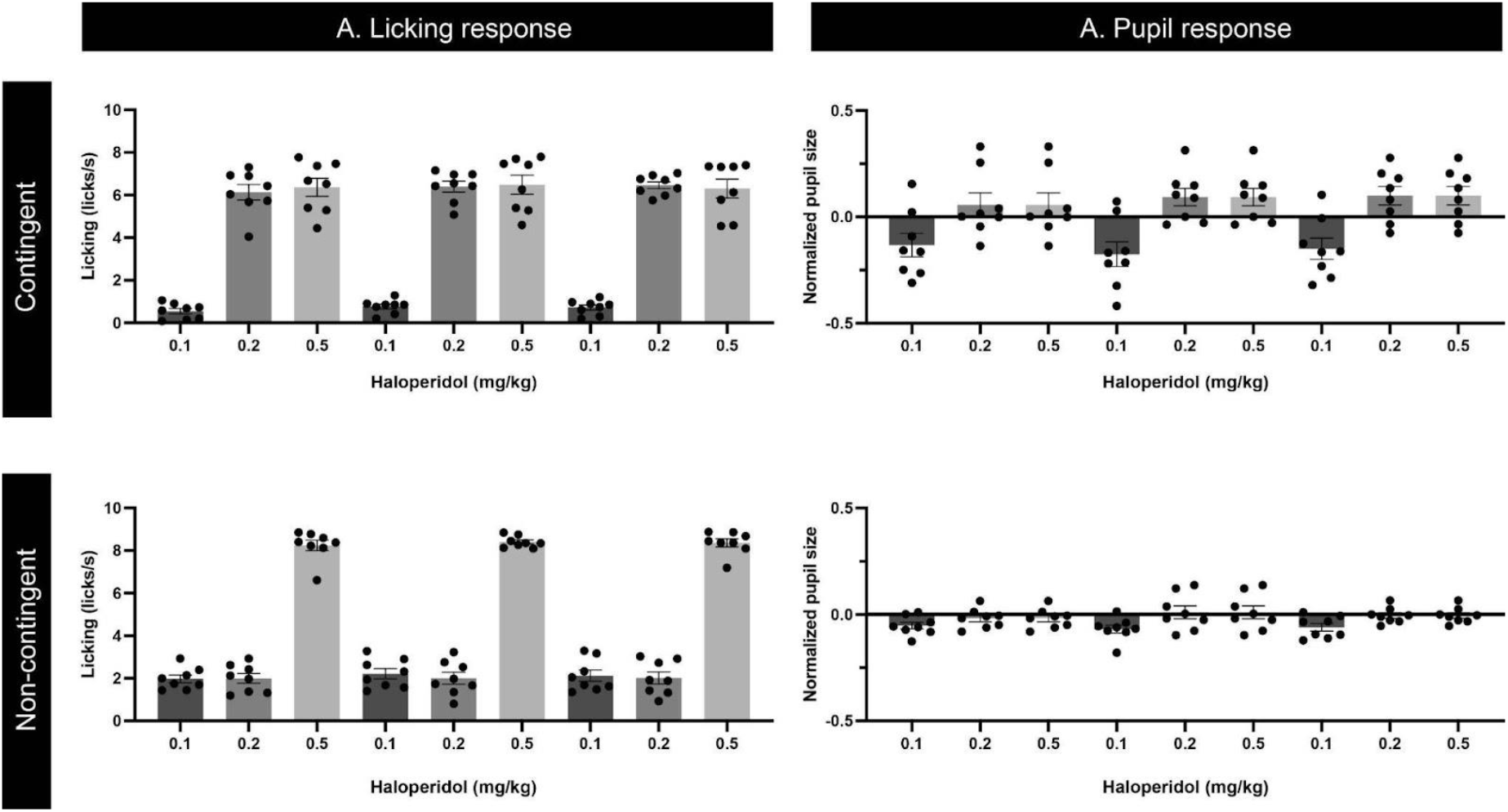
The amount of licking and pupil size at the time windows. The amount of licking and pupil size in Pre-CS, CS, and US periods of saline condition were separately shown by the previous dose of haloperidol injection.

## Notes

### Competing Interest Statement

The authors have declared no competing interest.

### Summary of Updates

Discussion and statistics in Results were mainly revised.

